# Weak influence of light and time of day on transcriptional response to SAV3 infection in Atlantic salmon (*Salmo salar*) smolts

**DOI:** 10.1101/2024.06.14.599015

**Authors:** Therese Solberg, Chandra S Ravuri, Mattis Jayme van Dalum, Jo Espen Tau Strand, David G Hazlerigg, Eva Stina Edholm, Alexander C West

## Abstract

Seventeen percent of aquaculture Atlantic salmon in Norway die after transfer from freshwater to seawater facilities, many from infectious disease. The degree to which these deaths are attributable to unseen weakness in smolt preparation is therefore an important concern. Strong evidence in mammals and supporting evidence in other teleosts show that the light environment shapes the immune response via entrainment of the circadian clock. These data are alarming as most smoltification protocols use constant light, which does not entrain the circadian clock and is associated with negative survival outcomes to disease challenge in other species. The goal of our study was to establish how different light environments affected the anti-viral immunity of Atlantic salmon smolts. To achieve this goal, we measured the transcriptional response of select type I and type II interferons and major histocompatibility complexes in the heart and head kidney following an intraperitoneal SAV3 viral challenge delivered under different lighting conditions. We began with two hypotheses. First, that compared with a regular light-dark cycle (LP), constant light (LL) would have a weaker resistance to viral challenge in Atlantic salmon smolts. Second, that Atlantic salmon smolts kept in a regular light-dark cycle would have altered immune sensitivity between the light and dark phases. To test our first hypothesis we photostimulated smoltification in parr through exposure to either LP or LL conditions for 6 weeks then challenged them by SAV3 infection. LP and LL groups showed indistinguishable measurements in classical smolt characteristics and comparable levels of SAV3 mRNA in the heart after 8 and 14 days post infection. They also showed comparable transcriptional responses of select type I interferons (*IFNa*, *IFNb*, *IFNc*) and both class I and class II major histocompatibility complexes (*MHC I* and *MHC II*) in both heart and head kidney tissue. *IFNγ* induction in the heart, however, was marked higher in the LL compared to the LP group. While this data suggests a heightened sensitivity or better immune response of the LL group to viral challenge, the consistent induction of *MHC II*, a key target gene of *IFNγ* signaling, to SAV3 infection in both LL and LP groups indicates that photoperiod may only have a minor impact on interferon regulation, rather than a more general influence on the viral immune response. To test our second hypothesis, we compared the transcriptomic response in the heart to SAV3 infection delivered either at the mid-light or mid-dark phase in the LP group. Both mid-light and mid-dark groups showed an upregulation of immune related genes and down-regulation of structural genes of mitochondria, but an analysis of the interaction between infection and time of day identified no differentially regulated genes. Overall, our data suggest that, unlike mammals, daily timing of viral infection may not play a major role in the immune response of Atlantic salmon.

**Highlights:**

- Atlantic salmon smolts die in large numbers following sea-water transfer in aquaculture
- Evidence in other species suggest that the light environment used in aquaculture contributes to these deaths
- We find that *IFNγ* and *MHC I* induction to a SAV3 infection in the heart is affected by photoperiod
- RNAseq analysis of hearts collected 14 days after PBS or SAV3 treatment, showed no statistical difference in SAV3-mediated gene induction between the light or dark phase.
- Our data suggests that, unlike in mammals, the viral immune response in Atlantic salmon is not affected by the time of day of infection.

## 1. Introduction

In early life, Atlantic salmon eggs hatch in freshwater and develop into a morph known as a parr. Once parr reach a critical size threshold they undertake a seasonally timed developmental transition into smolts; a morph that migrates from freshwater into the sea. The parr-smolt transition, known as smoltification, prepares Atlantic salmon for the new challenges of the marine environment (Stefansson et al., 2008). Smoltification, for example, alters gill physiology to meet the osmotic demands of seawater (Evans et al., 2005). Seawater migration, is however, also a major immunological stressor. Entry into the ocean environment exposes Atlantic salmon to a barrage of novel pathogens, and recent evidence suggests that smoltification involves significant immune remodeling of mucosal barriers that may be important for combatting the immune assault of seawater migration (Johansson et al., 2016a; Johansen et al., 2016b, Nuñez-Ortiz et al., 2018; West et al., 2021).

In 2023, 63 million Atlantic salmon died in Norwegian aquaculture following their transfer to sea cages (Fish Health Report 2023). Many of these fish died from pancreatic disease (PD), a condition characterized by pancreatic necrosis and severe heart inflammation; caused by the marine-dwelling salmonid alpha virus (SAV) (Deperasinska et al., 2018). In the interests of animal welfare and economic efficiency, it is important to determine whether rates of PD infection are due to a failure in the immune defenses of smolts, and in particular, if such failures derive from unseen weaknesses in aquaculture smoltification protocols.

In nature, the gradual change in annual photoperiod synchronizes Atlantic salmon smoltification (Duston and Saunders, 1990). Several weeks of exposure to short winter photoperiods sensitize the parr to longer photoperiods, so that the arrival of spring stimulates parr to develop into smolts (Björnsson et al., 1989; Strand et al., 2018). In contrast, aquaculture protocols commonly synchronize smoltification using abrupt shifts in extreme artificial photoperiods. Parr, raised in land-based facilities under constant light (LL), are exposed to short winter-like days for >6 weeks (SP; <12h light / 24h), before being returned to LL, which Atlantic salmon interpret as a long summer-like photoperiod (Duston and Saunders, 1990; Strand et al., 2018; Thrush et al., 1994; Handeland et al., 2001). Importantly, strong evidence in mammals and consistent evidence in fish suggest that the light environment used in aquaculture smoltification may have an under-appreciated impact on immune function (Scheiermann et al., 2018; Sacksteder et al., 2022).

In mammals, a regular 24h light-dark cycle is important for the entrainment of the endogenous circadian rhythm, which in turn coordinates daily cycles in the immune system (Scheiermann et al., 2018). Mice (Mus musculus), synchronized to a light-dark cycle, have radically different responses to an immune challenge depending on the time of day. The release of cytokines from stimulated macrophages, for example, is greater during the evening compared to the morning (Gibbs et al., 2011; Keller et al., 2009). Furthermore, intranasal viral infection of mice is greater during the inactive compared to the active phase (Edgar et al., 2016), showing that disease outcomes also change according to infection time. Significantly, both jet lag-like experiments and constant light lead to poorer health outcomes after viral challenge (Ehlers et al., 2018; Mizutani et al., 2017), underscoring the importance of a synchronized lighting environment for robust immunity.

We understand much less about the relationship between light, the circadian clock and the immune system in fish. Recent data, however, suggest an analogous relationship to that described in mammals (Sacksted et al., 2022; Du et al., 2017). The concentration of circulating cytokines in Japanese Medaka (*Oryzias latipes*), for example, are higher when treated with lipopolysaccharide (LPS; a surface membrane component present in most gram-negative bacteria) during the dark phase compared to the light phase, and appear to be linked to the expression of key circadian transcriptional activators (Onoue et al., 2019). Furthermore, rainbow trout (*Oncorhynchus mykiss*) exposed to constant light are less able to clear a louse infection (*Argulus foliaceus*) compared to fish housed under a regular 24h light-dark cycle (Ellison et al., 2021). These results support a conserved link between the circadian clock, lighting environment and immune function in fish. However, no study to date has explored this relationship in Atlantic salmon: a particularly significant omission given the standard use of LL in aquaculture smolt production.

The innate antiviral immune defense is, in large part, directed by interferons: a conserved group of signaling cytokines that are induced rapidly in response to viral nucleic acids (Weber, 2021; Dahle and Jørgensen, 2019). In mammals, interferons are divided into three families, type I (including *IFNα* and *IFNβ*), type II (*IFNγ*) and type III (*IFNλ*) (Robertsen, 2006). Type I and III interferons are produced by almost all cells in acute response to viral infection. Type II interferons, in contrast, are produced exclusively by cells of the immune system (Lee and Ashkar, 2018). Type II interferons are key activators of macrophages which are essential for the destruction of viral pathogens (Schroder et al., 2004). Interferon proteins ligate with interferon type-specific membrane bound cytokine receptors which in turn stimulate cells to enter an antiviral state (Negishi et al., 2018; Ivashkiv and Donlin, 2014). Interferon type specific receptors interact with different permutations of the JAK/STAT signaling pathway and interact with distinct transcription factor binding sites to elicit interferon type specific regulons (Negishi et al., 2018). Of chief importance among the suite of interferon-stimulated genes are the major histocompatibility complex (*MHC*) genes. *MHCs* are divided into *MHC class I* and *MHC class II* genes. Evidence suggests that *MHC class I* genes are primarily regulated by type II interferons whereas *MHC class II* genes are controlled by both type I and II interferons (Lee and Ashkar, 2018; Steimle et al., 1994). Once induced, MHCs are important for antigen presentation on the cell surface which stimulates downstream cell mediated responses which are key for eradication of virally infected cells.

Type I and II interferons are conserved between mammals and salmonids (Secombes and Zou, 2017). The interferon complement is more complex in salmon, however, in part due to the successive whole genome duplication events experienced by the salmonid group. For example, six different classes of type I interferons (*IFNa*, *IFNb*, *IFNc*, *IFNd*, *IFNe*, and *IFNf*) and two classes of type II interferons (*IFNγ-1* and *IFNγ-2*) have been identified in Atlantic salmon. Conversely, type III interferons (*IFNλ*) have not been identified in body fish suggesting a either a loss or later appearance of these genes in vertebrate evolution (Secombes and Zou, 2017). Although the evolution of the interferon system has followed different paths in salmonids and mammals, the major paths of induction (viral nucleic acids) and action (JAK/STAT pathway) appear to be conserved (Robertsen, 2018). In Atlantic salmon the classical MHC class I is represented by a single gene (*Ssa-UBA*). The MHC class II group in salmonids is represented by five genes in three major subgroups (referred to as DE, DB and DA). Only the DA group, however, displays classical characteristics of antigen presentation and is stimulated *in vitro* by recombinant IFNγ signaling (Dijkstra et al., 2013; Grimholt et al., 2020; Morales-Lange et al., 2021).

SAV3 infection stimulates the production of type I interferons *IFNa*, *IFNb* and *IFNc* in Atlantic salmon (Chang et al., 2016). Exogenous treatment with plasmids containing the *IFNa*, *IFNb*, and *IFNc* coding sequences, furthermore, is capable of protecting salmon from SAV3 infection for up to 10 weeks. In vitro work has also shown an antiviral role for *IFNγ* in response to SAV3, which may be partially mediated by *IFNa* induction (Sun et al., 2011). Together these data suggest that the antiviral response to SAV3 in Atlantic salmon is, in part, mediated through these subset of type I and II interferons.

The present study explores two hypotheses. First, that Atlantic salmon smolts housed under constant light would have an overall weaker immune response compared to Atlantic salmon house under a regular light-dark cycle. Second, that Atlantic salmon smolts housed under a light-dark cycle would show diurnal variation in their anti-viral immune response. To test our first hypothesis (LP vs LL), we stimulated smoltification under either LL or LP conditions and compared the immune response of the fish to intraperitoneal SAV3 challenge by quantifying the induction of class I and class II interferons with known antiviral activity against SAV3 (*IFNa*, *IFNb*, *IFNc* and *IFNγ*) and MHC class I and II transcripts. Against our expectations, the lighting environment had no effect on the rate and level of SAV3 infection and had only small effects on the transcription of *IFNa*, *IFNb*, *IFNc*, *MHC I* and *MHC II* genes in both heart and head kidney tissues. The most notable feature of our dataset was the induction of *IFNγ*, which was stronger under LL rather than LP in the heart. To test our second hypothesis (mid-light vs mid-dark), we compared the heart transcriptome of fish infected with SAV3 either in the middle of the light or dark phase. Although the RNAseq analysis identified a substantial transcriptional induction of immune-related genes and suppression of mitochondrial structural genes in response to SAV3 treatment compared to PBS controls, there were no significant difference between the heart samples infected in the mid-light group compared to the mid-dark group.

Although our study does not extend to defining disease outcomes of SAV3 challenge given at under different chronic lighting conditions or at mid-light and mid-dark timepoints, our data suggest that the robust link between circadian rhythms and the immune response to viral challenge in mammals is not a major feature of the Atlantic salmon immune response.

## 2. Materials and Methods

### 2.1 Ethical statement

The experiment was conducted in accordance with Norwegian and European legislations guidelines for animal research. The experiment protocol was approved by the Norwegian food safety authority (Mattilsynet FOTS 27998) in the fall of 2021.

### 2.2 Animal Husbandry

Atlantic salmon (Salmo salar, Aquagene commercial strain) were hatched and raised in freshwater under continuous light (LL; > 200 lux at water surface) with ambient temperature (∼10oC) at Havbruksstasjonen in Tromsø, Kårvik. Juvenile Atlantic salmon were kept in 500 L tanks and fed continually with normal pellet salmon feed (Skretting, Stavanger, Norway).

### 2.3 Experimental design *in vivo*

#### 2.3.1 Smoltification protocol

500 Atlantic salmon with a minimum body mass of 20 g were put in a common 100 L tank under LL. After 2 weeks of acclimation the Atlantic salmon were exposed to a winter photoperiod (short photoperiod; SP; 6h light / 24h) for 6 weeks. The cohort was then divided into two groups under different light regimes, one went through 6 weeks of constant light (LL group), and the other went through 6 weeks of long photoperiod (LP; 18h light / 24h). Both groups were kept in duplicate tanks with a minimum temperature of 10°C (See Figure 1A for experimental design).

**Figure 1:**
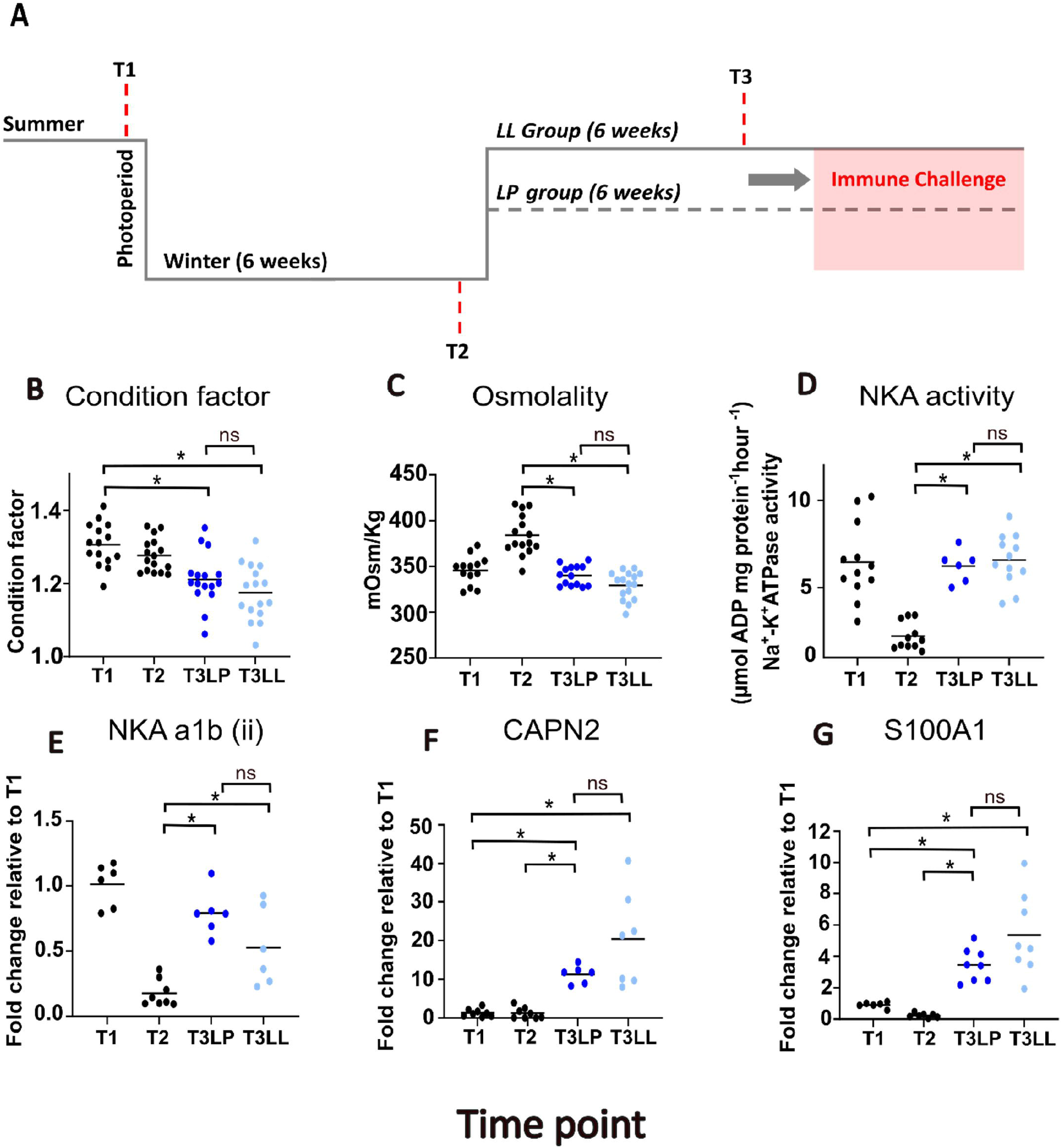
Experimental setup (A) Where the line represent the change in photoperiod, collection points are marked with T1-T3. (B) condition factor of salmon collected from different timepoints and lighting treatments (C) Blood plasma osmolality (mOsm/kg) following 24h seawater challenge (D) gill Na^+^ K^+^ ATPase activity (µmol ADP mg protein^-1^ hour^-1^) (E-G) qPCR values for the smolt marker genes *NKA a1b (ii)*, *CAPN2* and *S100A1*, respectively. Significantly different groups are marked with (*). There was no significant difference between T3 LP and T3 LL for any of the markers (marked with ns).

At experimental day 1 (Timepoint 1; T1), prior to the winter photoperiod under LL, 16 fish were randomly netted out from the tank. The fish overdosed with benzocaine (120 ppm) until unresponsive to external stimuli. Weight (g) and fork length were measured, and gill tissue was taken and snap frozen on dry ice. A gill filament sample was taken and stored in 100 µl of SEI buffer (26.67 g sucrose, 1.86 g Na_2_EDTA, 1.7 g imidazole diluted in 475 ml distilled water, pH: 7.3) for Na+, K+ ATPase (NKA) activity assessment. Blood samples were taken from the caudal vein of sea water challenge (SWC) fish using BD heparin vacutainers (Ref: 368494) and 27-gauge needles. Blood samples were centrifuged at 400xg for 10 min then plasma aliquots taken for osmolality measurement. All samples were stored at -80°C. The sampling was repeated on experimental day 42 (Timepoint 2; T2) at the end of the winter photoperiod, and experimental day 63 (Timepoint 3; T3) during LP and LL. Each timepoint also included a cohort of 16 fish which were exposed to full strength sea water (SWC) for 24h before overdose with benzocaine, blood sampling and plasma storage.

#### 2.3.2 SAV3 Immune challenge

After 6 weeks of winter photoperiod and 6 weeks of long photoperiod (LP or LL), 192 Atlantic salmon from the cohort were transferred to 8 circular seawater tanks with 24 fish in each tank. Four of the tanks contained LL smoltified fish and the other 4 contained LP smoltified fish. The two groups were kept in different rooms, still under their designated photoperiods. On experimental day 60, 2 of 4 tanks in both groups were intraperitoneally infected with 100 µl Salmonid alphavirus subtype 3 (SAV3) (1 × 105 TCID50). The other 2 tanks were intraperitoneally injected with 100 µl of PBS. SAV3 (PDV-H10-PA3) was originally provided by Professor Øystein Evensen, Norwegian University of life Sciences. The virus was propagated in CHH-1 cells, derived from heart tissue of a juvenile chum salmon (Onchorhynchus keta), in L-15 medium with 100 U/mL penicillin, 100 μg/mL streptomycin, and 5% FBS at 15°C. Virus titer was determined by the TCID50 method as described in (Strandskog et al., 2011).

For infection: fish were first aneasthetised in benzocaine (60 ppm) then the injection was administered on the midline, ∼1 pelvic fin length in front of the pelvic fin. In each tank, 12 fish were infected during the mid-dark phase of the LP group. The fish infected at were marked with a tattoo. The other 12 untattooed fish in each tank were infected 12 hours later during the mid-light phase (See Figure 3A for experimental design).

Heart and head kidney tissue samples were taken 8 days past infection and 14 days past infection. Samples were collected from 8 fish from each tank, 4 tattooed and 4 untattooed. The samples were snap frozen on dry ice and stored at -80°C until analyzed.

### 2.4 Analysis

#### 2.4.1 Osmolality and NKA assay

To test if the Atlantic salmon had smoltified properly in both LL and LP group we assessed their ability to hypo-omoregulate. Osmolality measurements of plasma samples from SWC fish (T1-T3) was conducted using the osmometer OSMOMAT® 030 GONOTEC. The osmometer was calibrated using 50 µl GONOTEC 850 calibration standard (Ref: 30.9.0850). The change in enzymatic activity of Na+, K+-ATPase was measured by NKA assay of the gill filaments following the method detailed in (Mccormick et al., 1993).

#### 2.4.2 RNA extraction and cDNA conversion

RNA was extracted from gill, head kidney tissues and leukocyte cells by using QIAgen RNeasy® plus mini kit (ThermoFisher, Ref:74134). RNA from hearts were extracted using QIAgen RNeasy® Universal mini kit (ThermoFisher, Ref: 73404). All RNA concentrations and were determined by using NanoDrop™ 2000/2000c. For tissue samples, 2000 ng RNA was reverse transcribed into cDNA using high-capacity RNA-to-cDNA kit (Applied biosystems™, Thermo Fisher, Ref: 4387406) following the manufacturer protocol.

#### 2.4.3 qPCR

The relative expression of smolt marker genes NKA a1b (II), S100A1 and CAPN2 were assessed by qPCR. So were the SAV3 marker non-structural protein1 (*Nsp1*), and immune marker genes for interferon Type 1, *IFNa-c*, Interferon type 2, *IFNγ*, and MHC class I and II. The primer sequences for the smolt markers were the same as used in (Iversen et al., 2020) and the immune marker sequences were obtained from (Svenning et al., 2019). The cDNA was amplified in Hard-Shell Low-Profile Thin-Wall 96-well skirted PCR Plates (Bio-Rad, Ref: HSP9601) containing 1 µl of cDNA template, 1µl forward primer, 1µl reverse primer (10nmol; table 1), 7µl nuclease-free water and 10 µl of SSo advanced universal SYBR Green qPCR kit (Bio-Rad, Ref: 1725272) for gill derived samples, and 10 µl Fast SYBR™ Green master mix (ThermoFisher, Ref: 4385612) as the reagent for heart and head kidney derived samples. The plate was read using the CFX96 Real-Time PCR Detection system (Bio-Rad) and the CFX Manager 3,1 software. The thermal cycling conditions were as follows: Gill samples. 10 minutes at 50°C and 5 minutes at 95°C followed by 40 cycles of 10 seconds at 95°C and 30 seconds extension phase. See table 1 for list of primer sequences and annealing temperature. Heart and head kidney. 2 min of 95°C followed by 40 cycles of 15 seconds at 95°C and 1 minute at 60°C. Relative expression was calculated using the ddCT method (Livak et al., 2001), where EF1-alpha was used as control for smolt markers, and the geometric mean of EF1-alpha (b) and (c) where used as control for Immune gene markers.

**Table 1:**
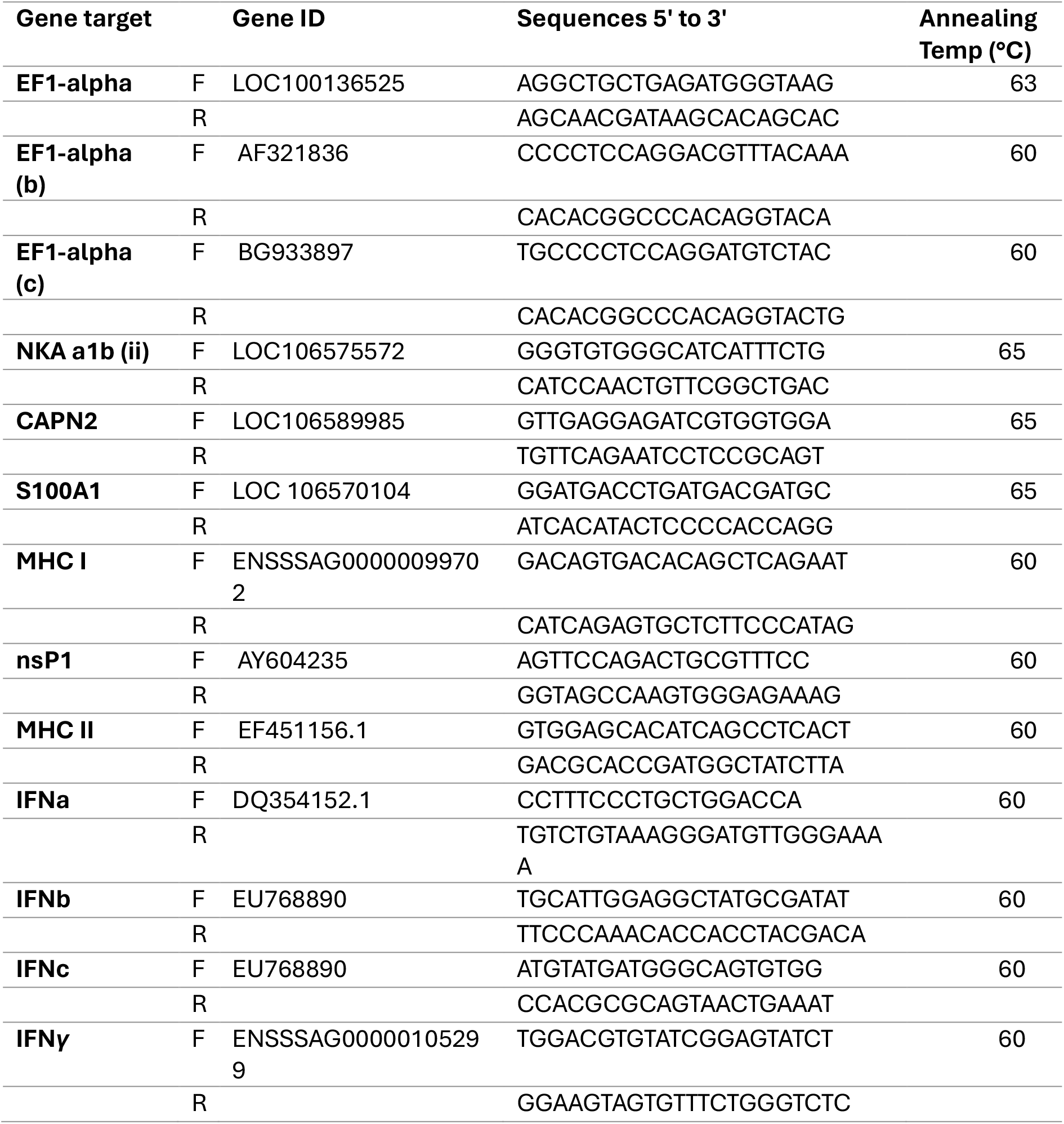
Primers used for qPCR.

#### 2.4.4 RNAseq

##### 2.4.4.1 Sample preparation, library preparation and sequencing

RNAseq was performed on heart samples from the LP group obtained 14 days past infection, only SAV3 treated samples which were above the infection threshold were used (see section 2.5) RNA quality was assessed using a tapestation (Agilant), all samples had an RNA integritiy number (RIN) >7. Library preparation and sequencing was performed by BGI (Hong Kong) who provided 20 million clean pair-end reads (PE100) using the DNBseq platform.

##### 2.4.4.2 RNAseq alignment

Prior to transcript quantification, adaptor sequences were trimmed off using fastp 0.23.2. Transcripts from single and paired-end RNAseq data were quantified based on the Atlantic salmon reference transcriptome version 3 (Ssal_v3.1, GCA_905237065.2) by using Salmon (v 1.1.0.) on mapping-mode. Library size was set to automatic detection, and we included the --gcBias flag which corrects for fragment-level GC biases in the input data. Bootstraps was set to 100. We loaded the salmon annotation file (.gff3) into R by using the package ‘ape’. Output files from Salmon containing read count estimates for each transcript were processed further in R studio (v 2023.6.0) and the package tidyR.

#### 2.5.2 Differential expression analysis

Analysis of differential transcript expression was conducted in Rstudio using the package edgeR (ver 3.42.4). Output files from Salmon containing estimated read counts for each transcript and bootstraps 100 were read by the edgeR function ‘catchSalmon’. To account for mapping uncertainty, transcript counts was divided by overdispersion. Overdispersion was automatically generated upon reading in the data as it was based on bootstraps data. Prior to analysis of differential expression, transcript counts were filtered by the function ‘filterByExpr’, setting an expression threshold of a minimum 10 counts per million (cpm) in a minimum number of samples; the treatment group with fewest samples.

Libraries of the transcript counts where scaled by the function ‘calcNormFactors’ which applies trimmed means of M-values (TMM) scaling. A multidimensional scaling plot (MDS plot) revealed one sample outlier in the PBS mid-dark group which was removed from analysis. Dispersion was calculated by the Cox-Reid profile-adjusted likelihood (CR) method using the function ‘estimateDisp’ (See edge R manual page 22 for ref). To find genes that that was differentially expressed we used a generalized linear model (GLM) approach, performing ‘glmQLFTest’ on our groups of comparison. Our contrast groups included: the interaction (SAV3 Treatment A - PBS treatment A) - (SAV3 treatment B - PBS treatment B), PBS treatment A - PBS treatment B, SAV3 treatment A - PBS treatment A and SAV3 treatment B - PBS treatment B. We used the package ‘DGEobj.utils’ to caluculate transcript per million (TPM). The test results were filtered by false discovery rate (FDR) with a threshold of 0.01. A subset from the TPM data was made for the immune markers from the qPCR analysis (*IFNa*, *IFNc*, *IFNγ* and *MHCs*) and TPM counts for each gene was combined.

Volcano plots (X axis: log2 fold change, Y axis: -log10 FDR) and MA plots (X axis: log2 mean expression, Y axis: log2 fold change), for the test results from SAV3 Treatment A - PBS treatment A and SAV3 treatment B - PBS treatment B, were generated in GraphPad prism.

#### 2.4.3 Gene ontology analysis

For gene ontology analysis, human genome nomenclature consortium identifiers from target gene lists were submitted to consensus path database (Kamburov et al., 2011). We used an over-representation analysis, normalised to a background of all the genes expressed in the dataset and exported gene ontology terms from levels 2-5 for biological process, molecular function and cellular compartment with a significance cut-off of p<0.01.

### 2.5 Data treatment and statistics

Condition factor was calculated according to the formula: Condition factor = (weight (g) × 100) / fork length3. All the data results were plotted in GraphPad Prim 9.0.0. Statistical analyses were performed in GraphPad and RStudio version 2022.02.0 (R Core team, 2020). To test if the data were normally distributed the residuals of the data was plotted in RStudio with the function qqnorm() followed by qqline(). The data was log transformed before statistical testing. To test if there were a significant difference in smoltification status between the groups (Groups: T1, T2, T3 LL and T3 LP) a post-hoc comparison was performed with the Tukey Honestly Significant Difference Method in RStudio. Infection success was evaluated by comparing nsP1 ct values for PBS and SAV3 injected groups. Based on this a threshold of ct 31 was made, where a value below was considered non-infected and therefore excluded from subsequent analysis. For the immune marker genes, a 3-way ANOVA in GraphPad was performed where P<0.05 was considered not infected.

## 3. Results

### 3.1 Smolt development is similar under constant light (LL) and long photoperiod (LP) conditions

We first compared classical smolt markers between the LL and LP raised smolts to detect baseline differences between the groups prior to SAV3 infection (See Figure 1A for experimental design). As expected, condition factor (an index of weight and length) decreased between T1 and T3 in both groups. There was, however, no significant difference in condition factor between the LL and LP groups (Figure 1B). We also measured several factors linked to the osmoregulatory capacity of the fish. Plasma osmolality of smolts exposed to 24h seawater challenge (SWC) was increased at T2 compared to T3, reflecting the typically poor hypo-osmoregulatory ability of fish under SP (Figure 1C). Similarly, we measured lower gill activity of Na+, K+ ATPase (NKA), a key ion pump believed to be essential for maintenance of osmotic balance, and lower gill abundance of NKAa1b (II), a smolt-associated subunit of the NKA holoenzyme (Mccormick et al., 2013), in T2 compared to T3 (Figure 1D and 1E). There were no significant differences in osmolality, NKA activity or NKAa1b (II) expression between the LL and LP groups. Finally, we measured the gill abundance or CAPN2 and S100a1, two genes whose transcript expression depends on prior exposure to winter-like photoperiods (Iversen et al., 2020). As anticipated, expression of these genes was low in T1 and T2 relative to the T3 groups. In line with our other findings, we measured no significant difference in the expression of CAPN2 and S100a1 between the LL and LP groups (Figure 1F and 1G). Overall, these data show that six weeks of LL and LP exposure produce comparable smolts, as characterized by classical smolt markers.

### 3.2 Induction of interferons and MHC class I in response to intraperitoneal SAV3 challenge are altered under constant light (LL) compared to long photoperiod (LP) conditions

We hypothesized that compared with a regular light-dark cycle (LP), constant light (LL) would have a negative impact on the induction of the early innate anti-viral immune response in Atlantic salmon smolts. To test our hypothesis we challenged both LL and LP groups with intraperitoneal infection of SAV3 virus and then measured the induction of select immune-associated genes 8 and 14 days post infection (dpi) in the heart, a major site of SAV3 infection (8), and head kidney, a primary/secondary lymphoid tissue in fish (Geven and Klaren, 2017). We chose to focus our characterization on the interferon response, which is key in innate anti-viral immunity, and represented here as the transcript induction of type I interferons (*IFNa*, *IFNb* and *IFNc*) and type II interferon (*IFNγ*). Further, we characterized the expression of the major histocompatibility complexes (MHC class I and MHC class II) which are essential for the processing and presentation of antigens and whose inducible expression is regulated by class I and class II interferons (Uribe et al., 2011).

We first determined the response to SAV3 challenge in the heart. Each SAV3-treated group had similar numbers of SAV3 positive fish and although we found that that the infection load, as inferred from detection of nsp1 transcript levels, measured by qPCR, was higher at 14dpi compared to 8dpi, there was no significant difference between the LP and LL groups (Supplementary Figure 1A). These data indicate that the overall infection rate in SAV3 challenged salmon was unaffected by light treatment. Analysis of infected fish revealed that *IFNa* was significantly induced at both 8dpi and 14 dpi with no difference between the LP and LL groups (Figure 2A). *IFNb* transcripts were below the level of detection by qPCR and *IFNc* levels were not significantly affected by the SAV3 challenge (Figure 2B). *IFNγ* induction, however, was stronger in the LL compared to the LP group, particularly evident at 14 dpi (Figure 2C). MHC class I transcripts were induced in both LP and LL groups at 14 dpi, however the induction was only evident at 8 dpi in the LP group (Figure 2D). Finally, we did not detect any change in MHC class II transcript abundance in either the LL or LP groups (Figure 2E).

**Figure 2:**
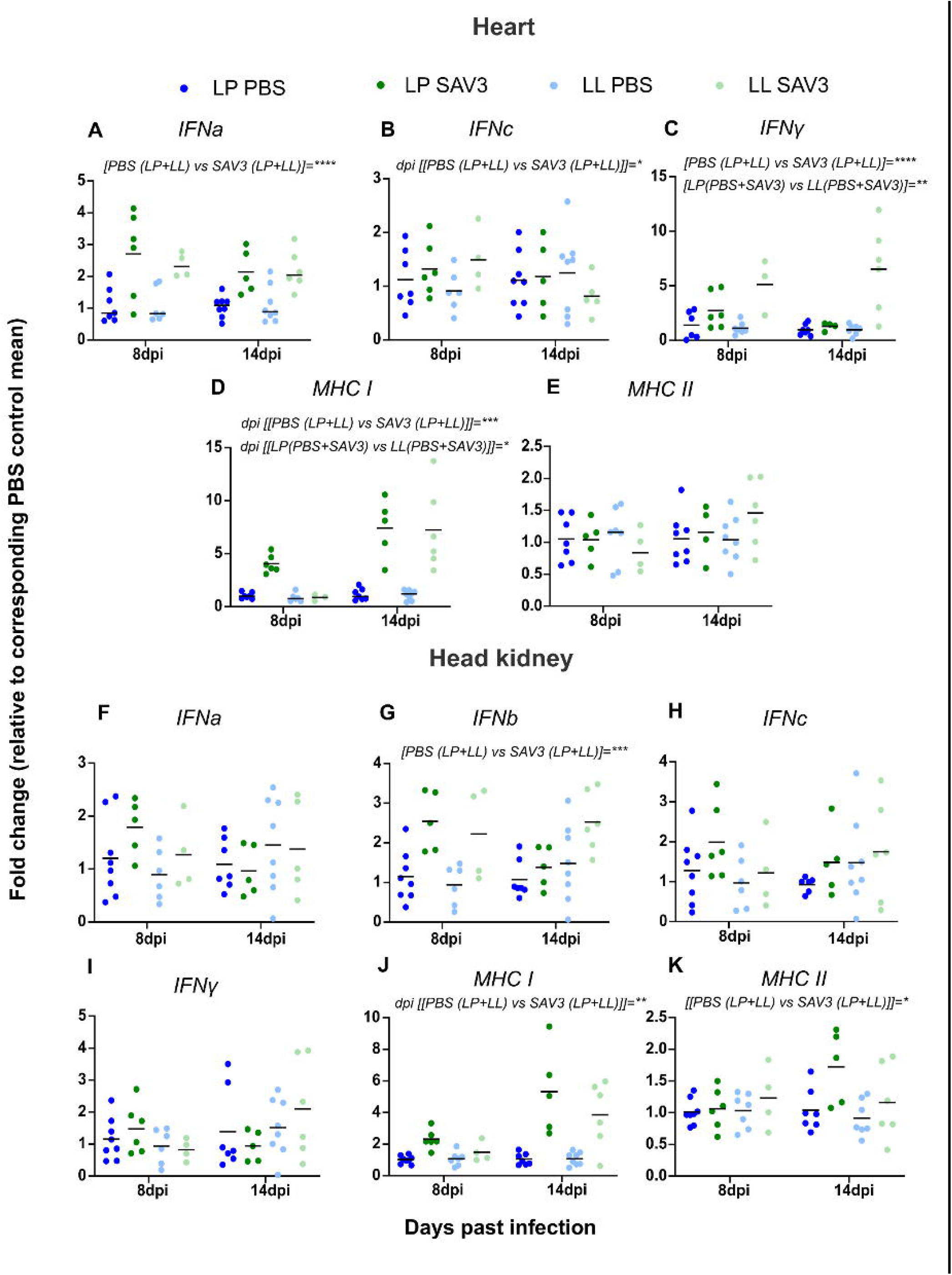
qPCR quantification of interferon and MHC class I and II in heart (A-E) and head kidney (F-K) collected at 8 and 14 days post infection (dpi). Data is presented as ddCT and significant differences following 3-way ANOVA detailed on each sub-panel.

We next characterized the response to SAV3 challenge in head kidney tissue. None of the samples had any detectable SAV3 virus, likely because the head kidney is not a site of infection for the SAV3 virus (Supplementary Figure 1B) (Desperasinska et al., 2018). We did not detect any increase or decrease in *IFNa*, *IFNc* or *IFNγ* transcripts in either the LL or LP groups (Figure 2F, 2H and 2I). We did however detect a modest increase in *IFNb* in both the LL and LP groups at both 8 and 14 dpi (Figure 2G). MHC class I was induced in both LL and LP groups at 14dpi, and MHC class II was weakly induced at 14dpi (Figure 2J and 2K).

Overall, our data shows that following SAV3 infection LL and LP housed fish exhibit similar rates of infection and transcriptional responses of a subset of interferons and MHCs, with the exception of *IFNγ* whose induction was higher in the heart under LL compared to LP conditions.

### 3.3 Transcriptome comparison of heart tissue from Atlantic salmon treated with SAV3 or PBS control from 14 days post infection showed strong evidence of SAV3-dependent immune response, but no evidence that mid-dark and mid-light infected fish had different immune responses

We hypothesized that Atlantic salmon smolts housed in a regular light-dark cycle would have altered viral immune response between the light and dark phases. To test this hypothesis, we challenged our LP group with an intraperitoneal infection of SAV3 or PBS control at the mid-light and mid-dark phases (ZT9 and ZT21, respectively) and then assessed their infection rates and anti-viral immune responses at 14 dpi from whole hearts (Figure 3A).

**Figure 3.**
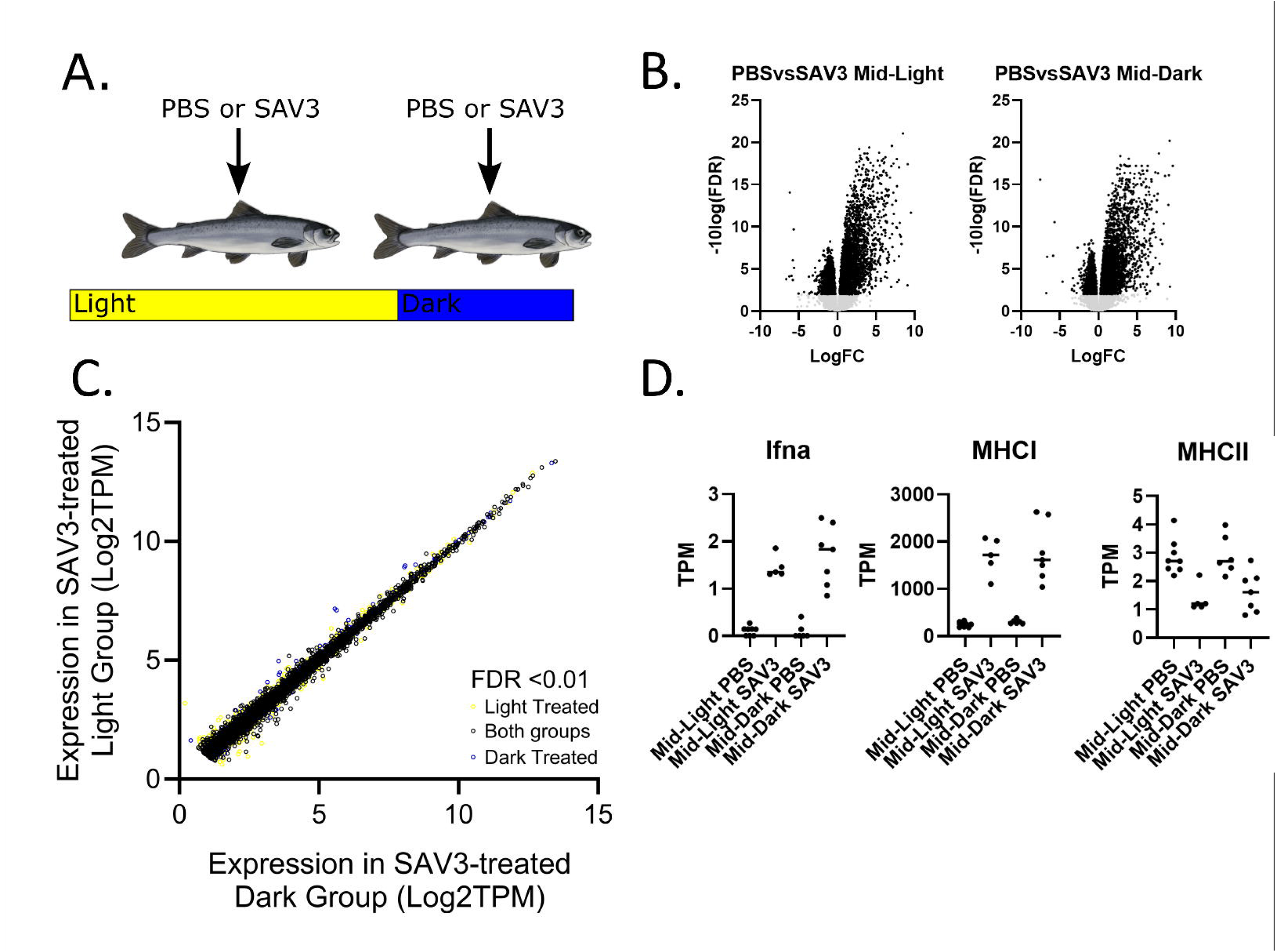
Transcriptomic analysis of heart tissue collected 14di after Atlantic salmon were injected with PBS or SAV3 at mid-light or mid-dark timepoints. A. Schematic representing the collection treatments and timepoints (see text for details). B. Volcano plot of differential expression of genes between PBS and SAV3 treated groups at mid-light and mid-dark timepoints. C. Scatter plot comparing the expression of SAV3-induced transcripts in mid-light and mid-dark groups. The plot shows genes that were significantly differentially expressed in the mid-dark (blue), mid-light (yellow) or both (black) groups with a significance cutoff of FDR <0.01 and absolute expression cutoff of 1 TPM. Transcript induction in response to SAV3 was not significantly different for any transcripts between mid-light and mid-light groups. D. Candidate gene expression.

The mRNA expression of the SAV3 nsp1 gene following infection was increased compared to PBS controls but not significantly different between mid-light and mid-dark groups, suggesting that, overall, the SAV3 infection was unaffected by light or time of day (Supplementary Figure 2). We next examined the response in the heart to SAV3 infection by comparing the transcriptomic profiles of PBS and SAV3-infected salmon from mid-light and mid-dark treatment groups (Figure 3B). Our analysis from the mid-light timepoint identified 8148 differentially regulated transcripts between the PBS and SAV3 groups, 4705 up-regulated and 3443 down-regulated (Figure 3B, Table S1). Comparison of the PBS and SAV3 groups in the mid-dark timepoint identified 7168 significantly different transcripts, 4216 up-regulated and 2952 down-regulated (Figure 3B, Table S2). Gene ontology analysis of the up-regulated gene sets identified categories related to immune function which were similarly over-represented in both the mid-light and mid-dark groups (Tables S1 and S2). Notably both mid-light and mid-dark groups were over-represented for: GO:0006955 Immune Response; GO:0045087 Innate Immune Response; GO:0009615 response to virus; GO:0042590 antigen processing and presentation of exogenous peptide antigen via MHC class I; GO:0060337 type I interferon signaling pathway; and GO:0060333 interferon-gamma-mediated signaling pathway. The down-regulated gene sets from both mid-light and mid-dark groups were associated with gene ontology groups related to mitochondrial structure and function (eg. GO:0044429 mitochondrial part, GO:0055114 oxidation-reduction process; Tables S1 and S2).

To compare the transcriptional responses between mid-light and mid-dark treated groups we performed an interaction analysis on our RNAseq datasets. In opposition to our hypothesis, our analysis identified no transcripts whose regulation by SAV3 are modified by the timing of infection (Table S3, Figure 3D). Examination of the dataset showed that *IFNb*, *IFNc* and *IFNγ* were all filtered out during the analysis due to their low abundance of mapped read counts. *IFNa*, *MHCI* and *MHCII* were evident in the analysed datasets and their response to SAV3 challenge was unchanged by the timing of infection (Figure 3E).

Taken together our data characterises a robust immune reaction in Atlantic salmon heart tissue following SAV3 infection. Our data, however, does not support our second hypothesis that immune sensitivity of Atlantic salmon smolts, entrained to a regular light-dark cycle, changes between the mid-light and mid-dark phases.

## 4. Discussion

Previous evidence from both mammals and fish support a role for the light environment in optimal immune defense (Scheiermann et al., 2018; Sacksteder et al., 2022). Our present study explores this relationship in Atlantic salmon during smoltification, a key physiological and environmental transition that is associated with substantial losses in aquaculture due to disease (Stefansson et al., 2008). Our data shows that the chronic light environment alters the transcriptional induction of *IFNγ* and MHC class I transcripts in the heart following SAV3 challenge. However, transcriptomic analysis comparing mid-light and mid-dark SAV3 challenged heart tissue showed no statistical interaction between infection and time of day.

As anticipated, both LL and LP groups showed reduction in condition factor following SP exposure. Fork length of the LL group was longer (18.6 cm, SD: 0.86) compared to the LP group (17.1, SD: 0.9) in agreement with prior work that reports higher growth rates in LL housed salmon compared to other ‘summer-like’ photoperiods (Kråkenes et al., 1991 and Strand et al., 2018). This is likely due to elevated circulating GH, a hormone which simulates food intake and fork-length growth, and whose release is proportional to the photostimulatory daylength (McCormick et al 1995, Stefansson et al 1991, JETS 2018, Johnsson and Bjornsson 1994). Condition factor was, however, indistinguishable between the LL and LP groups showing that the development of a marine body shape was conserved between the two chronic light treatments.

Blood plasma osmolality following a 24h seawater challenge was increased on transfer to SP then decreased again following transfer to either LP or LL groups. We also observed consistent reduction in NKA activity and NKA a1b (ii) gene expression in gill samples following transfer to SP which recovered upon transfer to either LL or LP. Together these data are consistent with a changing hypo-osmoregulatory capacity typical to the smoltification transition in Atlantic salmon (Iversen et al., 2020; Nilsen et al., 2007). We also measured the gill expression of winter-dependent genes *S100a1* and *Capn2* whose expression pattern was strongly stimulated following transfer to either LP or LL conditions following SP exposure, comparable with previous studies (Iversen et al., 2020, West et al., 2021). Importantly, each of these measures were indistinguishable between the LP and LL groups suggesting that these fish were in similar physiological states at the point of SAV3 infection.

Our first hypothesis was that compared to the LP group, the LL housed fish would show evidence of a weaker immune response to SAV3 challenge. We identified no SAV3 virus in the head kidney, likely as the head kidney is not a primary infection site of the SAV3 virus (Deperasińska et al., 2018). Induction of interferons in the head kidney was weak or absent, we did however, detect a strong *MHC I* signal (Figure 2). We ascribe this difference firstly to the low viral burden in the head kidney meaning that there is likely no direct interferon activity in the tissue, and secondly to the head kidney’s role as a secondary lymphoid organ, meaning that MHC I expressing cells are likely recruited to the tissue for antigen presentation (Press & Evensen, 1999). Crucially, in relation to our hypothesis, we found no evidence that chronic exposure to LL or LP influences the expression of our panel of interferons and *MHC* genes to SAV3 exposure.

SAV3 infection load in heart tissue was identical between LP and LL groups both at 8dpi and 14 dpi suggesting that the level and progression of SAV3 infection is similar between the two groups (Supplementary Figure 1). While we did find differences in the expression of *IFNγ* (stronger induction at 14 dpi in the LL group) and *MHC I* (no induction in LL group at 8 dpi), however, our data does not describe consistently weaker response in either the LP or LL groups. For example, IFNγ signaling is causally linked to the induction of both *MHC I* and *MHC II* in Atlantic salmon (Grimholt et al., 2020) and yet the stronger induction of *IFNγ* at 14 dpi in LL compared to LP does not stimulates a stronger induction in these key target genes. Taken together, in opposition to our first hypothesis, we do not observe a coordinated influence of chronic photoperiod exposure on the innate viral immune response.

Our second hypothesis was that Atlantic salmon would have an altered immune response to SAV3 virus delivered at mid-light and mid-dark timepoints. In striking contrast to our hypothesis, we identified no genes whose induction or repression following SAV3 challenge was altered depending on the time of day of infection (Figure 3C). These data may reflect differences in circadian coordination between Atlantic salmon and other vertebrates. Our recent work has explored how canonical clock genes are regulated in Atlantic salmon smolts. Under a daily light-dark cycle we identified many conserved clock genes cycled in expression in the optic tectum we found far fewer cyclic genes in the gill and saccus vasculosus. Crucially, while circadian rhythms in the transcriptional profiles of many of these genes persisted in the optic tectum, circadian expression of clock genes was abolished in the gill and only persisted on a single gene in the saccus vasculosus (West et al., 2018). These data contrast with work in zebrafish where all tissues are capable of their own light detection and self-sustaining circadian rhythmicity (Whitmore et al., 1998, Davis et al., 2015). The implication of these data in the context of our present study supports a narrative of weakened peripheral clocks in Atlantic salmon. Most of our knowledge of circadian immunology derives from human and rodent studies, groups who both have high amplitude circadian rhythms in their immune responses (Scheiermann et al., 2018). These groups also experience daily cycles in their exposure to novel pathogens due to their sleep-wake cycles (Tognini et al., 2017), which is distinct from the experience of Atlantic salmon who likely encounter less daily variation in pathogen exposure. We suggest, therefore, that the ultimate drivers of circadian regulation on mammalian human and rodent immune systems are likely to be stronger than those in Atlantic salmon and that this may have diminished the adaptive value of strong circadian regulation of the immune system in salmon.

Taken together our work suggests that, unlike mammals, light and time of day does not have a strong influence on the innate immune response in Atlantic salmon smolts. We note however, the Atlantic salmon immune system is likely dramatically different during different life history stages and urge caution in extrapolation of our data to other immune challenge paradigms.

## Acknowledgements

The authors thank all the animal staff at Kårvik havbruksstasjonen for their expert care of the research animals.

## 5. Contributions

Conceptualization: TS, DGH, ESE and ACW; Data generation: TS, CSR, JETS, ESE, ACW; Data analysis: TS, MJVD, JETS, ESE, ACW; Manuscript preparation: All; Funding: DGH.

---

Supplementary Figure 1. Expression of SAV3 mRNA marker qnsp1 at 8 and 14 days post infection (dpi) following PBS control or SAV3 treatment in heart and head kidney tissue. See text for details.

Supplementary Figure 2. Expression of SAV3 mRNA marker qnsp1 14 days post infection (dpi) following PBS control or SAV3 treatment at mid-light or mid-dark timepoints in heart tissue. Statistics: 2-way ANOVA. P****<0.001.

