## Supplementary figures and images for "Weak influence of light and time of day on transcriptional response to SAV3 infection in Atlantic salmon (*Salmo salar*) smolts"

### Supplementary Figure 1

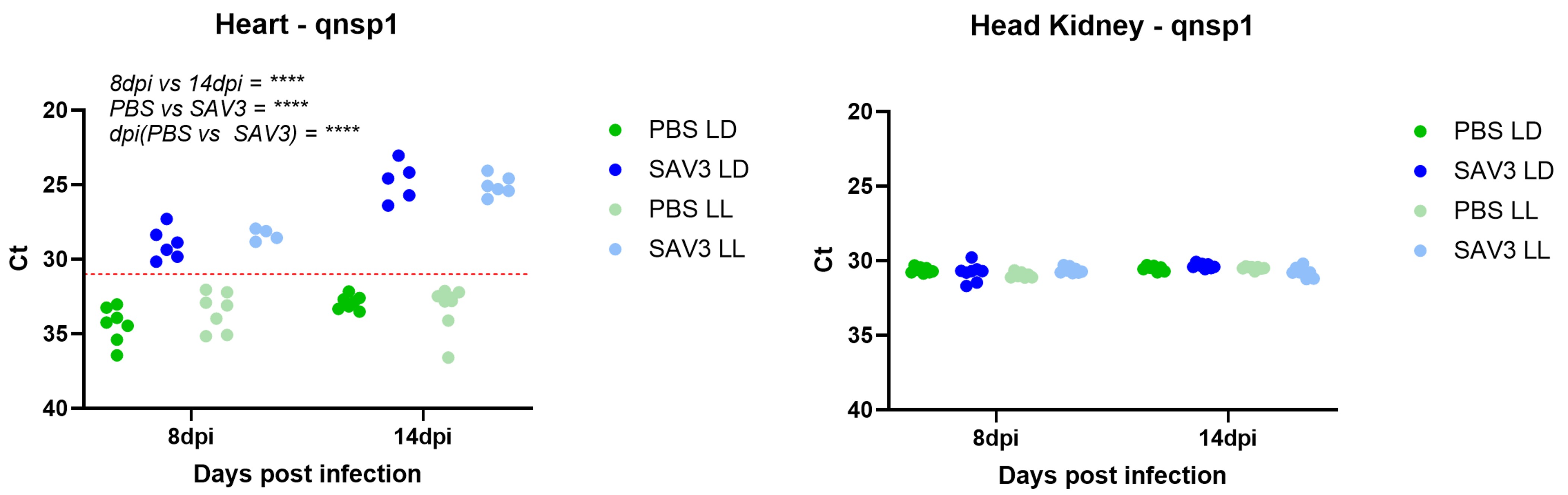

### Supplementary Figure 2

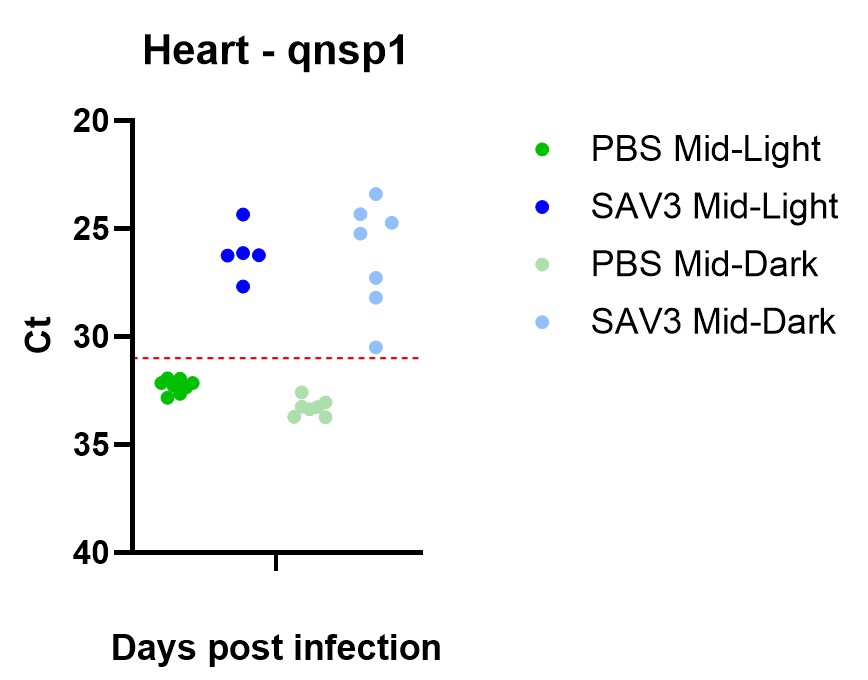
